# Early urinary candidate biomarkers and clinical outcomes of intervention in a rat model of experimental autoimmune encephalomyelitis

**DOI:** 10.1101/205294

**Authors:** Mindi Zhao, Yameng Zhang, Jiangqiang Wu, Xundou Li, Youhe Gao

**Author notes:** The authors contribute equally to the study (Mindi Zhao, Yameng Zhang, Jiangqiang Wu). To whom correspondence may be addressed. Email address: Youhe Gao.

## Abstract

Multiple sclerosis is a chronic autoimmune demyelinating disease of the central nervous system and is difficult to diagnose in early stages. Without homeostatic control, urine was reported to have the ability to accumulate early changes in the body. We expect that urinary proteome can reflect early changes in the nervous system. In this study, the early urinary proteome changes in a most employed multiple sclerosis rat model (experimental autoimmune encephalomyelitis (EAE)) were analyzed to explore early urinary candidate biomarkers, and early treatment of methylprednisolone were used to evaluate the therapeutic effect. Compare with controls, twenty-five urinary proteins were altered at day 7 when there were no clinical symptoms and no obvious histological changes. Among them, twenty-three have human homologs and fourteen were reported to be differently expressed in the serum/cerebrospinal fluid/brain tissues of multiple sclerosis patients or animal models. Functional analysis showed that the dysregulated proteins were associated with asparagine degradation, neuroinflammation and lipid metabolism. After the early treatment of methylprednisolone, the incidence of encephalomyelitis in the intervention group was only 1/13. This study demonstrates that urine may be a good source of biomarkers for the early detection of multiple sclerosis and early treatment can significantly delay disease progression. These findings may provide important information for early diagnosis and intervention of multiple sclerosis in the future.

## Background

Multiple sclerosis is a chronic autoimmune demyelinating disease of the central nervous system that are characterized by both inflammatory components and neurodegeneration [1]. It contributes to a significant proportion of neurologic morbidity in young adults [2], but the causes of multiple sclerosis are not fully understood. Magnetic resonance imaging (MRI) is the most common diagnostic tool for multiple sclerosis. Additionally, some clinical features may help to diagnose the disease. However, few clinical manifestations are specific to multiple sclerosis, and MRI lacks specificity for the early stages of the disease. In addition, when the clinical symptoms developed, the medical treatment is always not timely enough. And the early intervention of multiple sclerosis is still a matter of concern. Early treatment to slow or reverse inflammatory lesion formation is advocated as an effective way to prevent accumulation of disability [3].

Without homeostatic control, urine can reflect most changes in the body and thus can be a better source of biomarkers [4]. Numerous biomarkers of various diseases can be detected in urine, and some of these biomarkers perform even better than plasma biomarkers [5]. Advancements in mass spectrometry (MS) have made it possible to uncover new distinct molecular components. During the past few years, proteomics approaches have been used to investigate changes in urinary proteins/metabolites of multiple sclerosis patients. For example, proteins such as trefoil factor 3 and lysosome-associated membrane protein 2, both of which are related to immune responses, were differentially expressed in two phases (the third trimester of pregnancy and the postpartum period) of multiple sclerosis patients [6]. Multiple sclerosis shares many overlapping clinical features with neuromyelitis optica spectrum disorders, but the treatment strategies differ substantially for these two diseases, and thus several urinary proteomic/ metabolome studies have been conducted to differentiate these two diseases [7, 8].

Experimental autoimmune encephalomyelitis (EAE) is the most commonly employed model for multiple sclerosis [9], and it has been a powerful tool for studying relevant mechanisms in multiple sclerosis as well as for translating the findings into clinically meaningful therapeutic approaches [10]. The clinicopathologic characteristics of the EAE model, including inflammation and demyelination of the central nervous system, are like those of multiple sclerosis [11]. Thus, studies using the EAE model have provided new insights into the pathogenesis and pathophysiology of multiple sclerosis. Because multiple sclerosis is difficult to diagnose at early stages in clinical practice, in this study, the EAE model was used for the discovery of early urinary biomarkers of multiple sclerosis. And for animal models, it is convenient to limit confounding factors to understand the onset of diseases [12].

In the present study, we used two-dimensional high-resolution mass spectrometry to investigate changes in the urinary proteome during the early stages of multiple sclerosis in EAE rat models. On this basis, we explored the effect of early intervention with methylprednisolone. Urine samples were analyzed by liquid chromatography coupled with tandem mass spectrometry (LC-MS/MS) before the onset of disease symptoms (day 7). The changed proteins were then correlated to neurological functions by network and canonical pathway analysis. The study showed the early diagnosis and intervention of multiple sclerosis is important, and urine proteome could reflect early changes of EAE models.

## Methods

### Experimental rats

Fifty-six male Lewis rats (8 weeks old) were purchased from the Institute of Laboratory Animal Science, Chinese Academy of Medical Science & Peking Union Medical College. The experiment was approved by the Institute of Basic Medical Sciences Animal Ethics Committee, Peking Union Medical College (Animal Welfare Assurance Number: ACUC-A02-2014-007). The study was performed according to guidelines developed by the Institutional Animal Care and Use Committee of Peking Union Medical College.

### EAE rat models for early diagnosis

Thirty rats were randomly divided into two groups, namely, the EAE group (n=15) and the control group (n=15). EAE was induced in Lewis rats with myelin basic protein (MBP) as previously described [13]. Rats in the EAE group were immunized with subcutaneous injections of 100 µg MBP (Sigma) emulsified in 5 mg/mL complete Freund’s adjuvant (CFA) containing Mycobacterium butyricum (Sigma). Rats in the control group were administered with CFA and infused with an equal amount of saline. On day 7, all rats were placed in metabolic cages to collect urine. Three pairs of rats in the two groups were sacrificed at each timepoint (day0, 7, 14, and 21), and tissues were collected for histological analyses. Body weight and neurological impairment scores were evaluated daily. The progression of EAE was measured daily based on neurological impairment and scored from 0 to 5 as follows [14]: grade 0, no symptoms; 0.5, mild floppy tail; 1, floppy tail; 2, hindlimb weakness; 3, severe paraparesis; 4, tetraparesis; and 5, moribund.

### EAE rat models for methylprednisolone early intervention

Twenty-six rats were randomly divided into three groups, the early treatment group (n=13), the late treatment group (n=7), and the control group (n=6). EAE models were induced in all the rats as described before. According to the results of the EAE rat models for early diagnosis as above, the early treatment group received intraperitoneal injection of methylprednisolone (Pfizer) on the 9th day after modeling, and the late treatment group received intraperitoneal injection of methylprednisolone after disease progression (day10, 11 and 12). Methylprednisolone was injected at a dose of 30 mg/kg for 5 consecutive days. The control group was injected with the same amount of normal saline after onset. Body weight and neurological impairment scores were evaluated daily as mentioned above.

### Histological analysis

On days 7, 14 and 21, spinal cord samples were harvested from both EAE and control groups and dissected after blood withdrawal. After being fixed in 4% paraformaldehyde, the samples were embedded in paraffin. Sections of paraffin-embedded heart samples were stained with hematoxylin and eosin (H&E) for the evaluation of inflammatory foci.

### Urine sample preparation

After collection, urine was immediately centrifuged at 2,000 g to remove pellets. Urinary proteins were extracted by adding three volumes of acetone. After centrifugation, proteins were dissolved in lysis buffer (8 M urea, 2 M thiourea, 25 mM dithiothreitol and 50 mM Tris). The urinary proteins were then denatured with dithiothreitol, alkylated with iodoacetamide, and digested with trypsin (Promega) (1:50) at 37°C overnight using filter-aided sample preparation methods as previously described [12]. The digested peptides were desalted using Oasis HLB cartridges (Waters, USA).

Urine samples (from fifteen rats with EAE and fifteen control rats) collected on day 7 were used for MS analysis. As the tandem mass tag (TMT) reagents (Thermo Fisher Scientific) have six channels, random five samples were mixed to one sample in each group (total six group). Peptides in each sample were labeled with 126, 127, 128, 129, 130 and 131 according to the manufacturer’s instructions. The labeled peptides were mixed and then analyzed with two-dimensional liquid chromatography-MS/MS (LC-MS/MS).

### Reverse-phase liquid chromatography separation

TMT-labeled peptides were fractionated using offline high-pH reverse-phase liquid chromatography (RPLC) columns (XBridge, C18, 3.5 μm, 4.6 mm × 250 mm, Waters). Peptides were diluted in buffer A1 (10 mM NH_4_FA in H_2_O, pH = 10) and then loaded onto the RPLC column. The elution buffer consisted of 10 mM NH_4_FA in 90% acetonitrile (pH = 10; flow rate = 1 mL/min; 60 min). The eluted peptides were collected at a rate of one fraction per minute. After lyophilization, 60 dried fractions were resuspended in 0.1% formic acid and combined into 15 fractions; for example, fractions 1, 16, and 31 were combined with fraction 46.

### LC-MS/MS analysis

Each fraction was analyzed in duplicate using a reverse-phase C18 (3 μm, Dr. Maisch, Germany) self-packed capillary LC column (75 μm × 120 mm). The elution gradient was 5–30% buffer B (0.1% formic acid in acetonitrile: flow rate 0.3 μL/min) for 60 min. A TripleTOF 5600 MS system was used to analyze the eluted peptides. The MS data were acquired using data-dependent acquisition mode with the following parameters: 30 data-dependent MS/MS scans per full scan; acquisition of full scans at a resolution of 40,000, and acquisition of MS/MS scans at a resolution of 20,000; rolling collision energy; charge state screening (including precursors with +2 to +4 charge states); dynamic exclusion (exclusion duration 15 s); an MS/MS scan range of 250-1800 m/z; and a scan time of 50 ms.

### Data analysis

All MS/MS spectra were analyzed using the Mascot search engine (version 2.4.1, Matrix Science), and proteins were searched against the SwissProt_2014_07 database (taxonomy: Rattus, containing 7,906 sequences). Carbamidomethylation of cysteines was set as fixed modifications, the precursor mass tolerance and the fragment mass tolerance were set to 0.05 Da, and two missed trypsin cleavage sites were allowed. To obtain convincing results, proteins were filtered using the decoy database method in Scaffold (version 4.3.2, Proteome Software Inc., Portland). The false discovery rate (FDR) of proteins was set below 1.0%. Each protein contained at least two unique peptides. Scaffold Q+ software was employed for the quantification of TMT labeling. The statistical test used in Scaffold Q+ was permutation; the changed proteins were defined based on a fold change > 1.5 and a p value < 0.05.

### Functional analysis

For functional analysis, deregulated urinary proteins identified by MS were further annotated by Ingenuity Pathway Analysis (IPA) software. After importing the information on the deregulated proteins and their fold changes into the Ingenuity website, we analyzed the affected canonical pathway, networks, and related diseases. Biological functions assigned to each canonical pathway were ranked according to the significance of that biological function in the pathway.

## Results and discussion

### Workflow for quantitative proteomics analysis of EAE rats

To determine the urinary candidate biomarkers of EAE, we immunized rats with MBP to establish an EAE model. Body weight indexes, neurological impairment scores and histopathological characterization were used to evaluate disease progression. On day 7 after immunization, few rats with EAE display clinical symptoms, and this phenomenon was identified as the time point to study early candidate biomarkers. To illustrate the importance of early diagnosis, early intervention was administrated at the onset of symptoms on day 9 and late intervention was administrated as the disease progression in some rats. For comprehensive and comparative analyses, the urinary peptides were labeled with TMT. The TMT-labeled samples were separated into 15 fractions by offline RPLC, and each fraction was then analyzed in duplicate by LC-MS/MS. The workflow for the quantitative proteomics analysis is shown in Figure 1.

**Figure 1.**
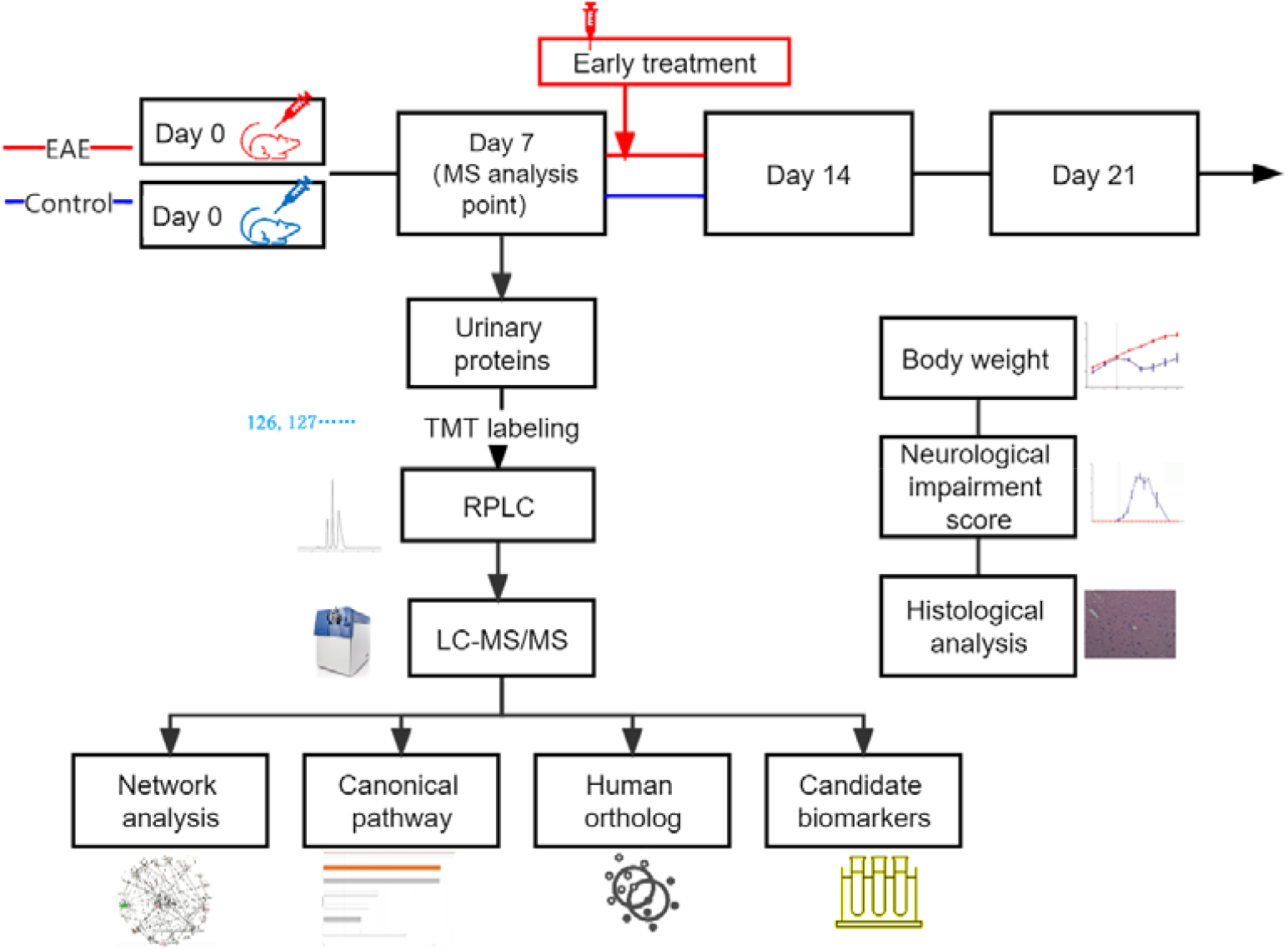
Workflow of the study. The study was divided to two parts, the early biomarkers discovery and the early intervention study. Urinary proteins at day 7 were identified by TMT labeling followed by LC-MS/MS. For the intervention study, EAE rats were received intraperitoneal injection of methylprednisolone on the 9th day after modeling in the early treatment group.

Raw data were searched against the SwissProt database for the taxonomy Rattus and then imported into Scaffold Q+ software for protein identification and quantification. If the identified peptides were shared between two proteins and could not be separated based on the MS data, the two proteins were grouped and annotated as one protein group. In total, 613 proteins that consisted of at least two peptides were identified at a 1% FDR at the protein level. Among these proteins, 566 high-confidence proteins were quantified by TMT labeling analysis in duplicate.

### Body weight and neurological impairment characterization

Parameters of the EAE and control groups were evaluated from day 0 to day 21. Both MBP-immunized Lewis rats and control rats exhibited a consistent 10 to 15% increase in body weight from the first day to the seventh day. In the first week, no significant difference in body weight was observed. From the eighth day onward, the weight of rats in the control group remained on the rise and were higher than that of rats with EAE. From day 11 onward, there was a marked decrease in the weight of the EAE group (225.1 ± 16.9 g), whereas the weight of the normal group was 268.1 ± 10.1 g. From day 13 to day 14, the weight of the EAE group stopped decreasing and remained unchanged. By day 21, the last time point examined in the study, the weight of the EAE group returned to the value on day 7. As shown in Figure 2A, the difference in body weight between the EAE and control groups was large, indicating possible impairment due to MBP immunization. Additionally, Lewis rats with EAE displayed a monophasic clinical course and spontaneous recovery, both of which are also consistent with previous results [15].

**Figure 2.**
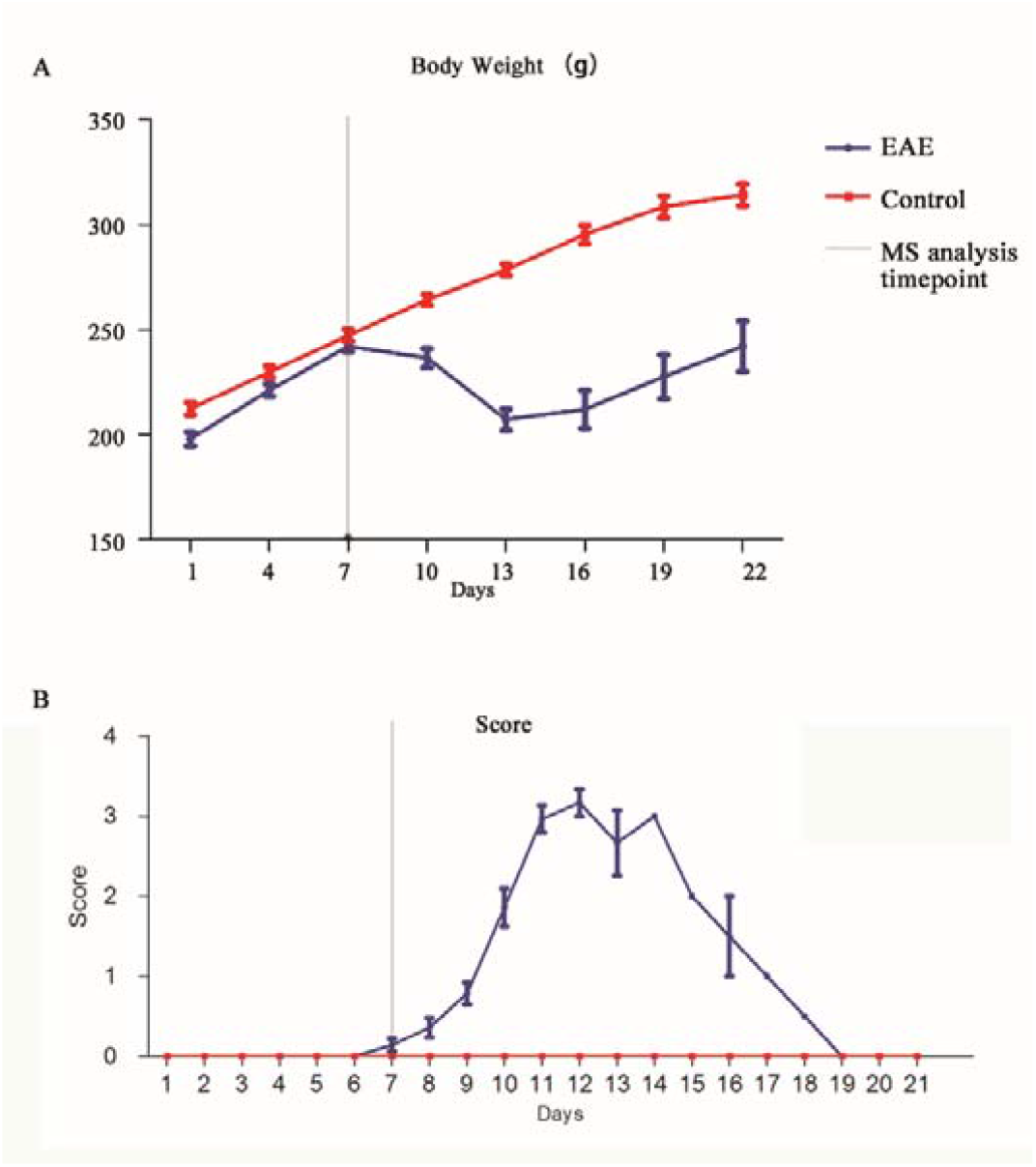
Changes in body weight and neurological damage score during the experiment. The red line indicates the control group, and the purple line indicates the EAE group.

In the present study, no rats died or reach “grade 5” in disease severity. From day 0 to day 7, consistent with the lack of negative changes in body weight, neither the MBP-immunized Lewis rats nor the control rats displayed any abnormal clinical symptoms. All rats immunized with MBP demonstrated typical clinical manifestations of neurological impairment lasting from 8 to 21 days post-immunization. On the eighth day, which is also the time when the weight began to decrease in rats with EAE, some of the MBP-immunized rats developed mildly flaccid tails, and this symptom became obvious in nearly all rats with EAE by the ninth day. From day 12 to day 14, the rats in the EAE group exhibited progressive bilateral hindlimb paralysis (grades 2 to 3), which was also the most severe symptom observed in this study. Then, the rats recovered from paralysis. By the end of the experiment, these effects disappeared (Figure 2B).

### Histopathological findings during the progression of EAE

Histopathological examinations were performed by H&E staining of spinal cord sections from rats with EAE to evaluate the severity of disease (Figure 3). During the early stages after initial immunization, very few infiltrated inflammatory cells were observed. On day 14, H&E staining revealed the presence of numerous inflammatory infiltrates in the parenchyma of the spinal cord and perivascular area. On day 21, also known as the recovery stage, some inflammatory lesions had disappeared; nevertheless, unlike the absence of inflammatory cell infiltration in the control samples, several inflammatory cells were still detected in the EAE samples. These pathological changes demonstrated the successful induction of EAE in Lewis rats.

**Figure 3.**
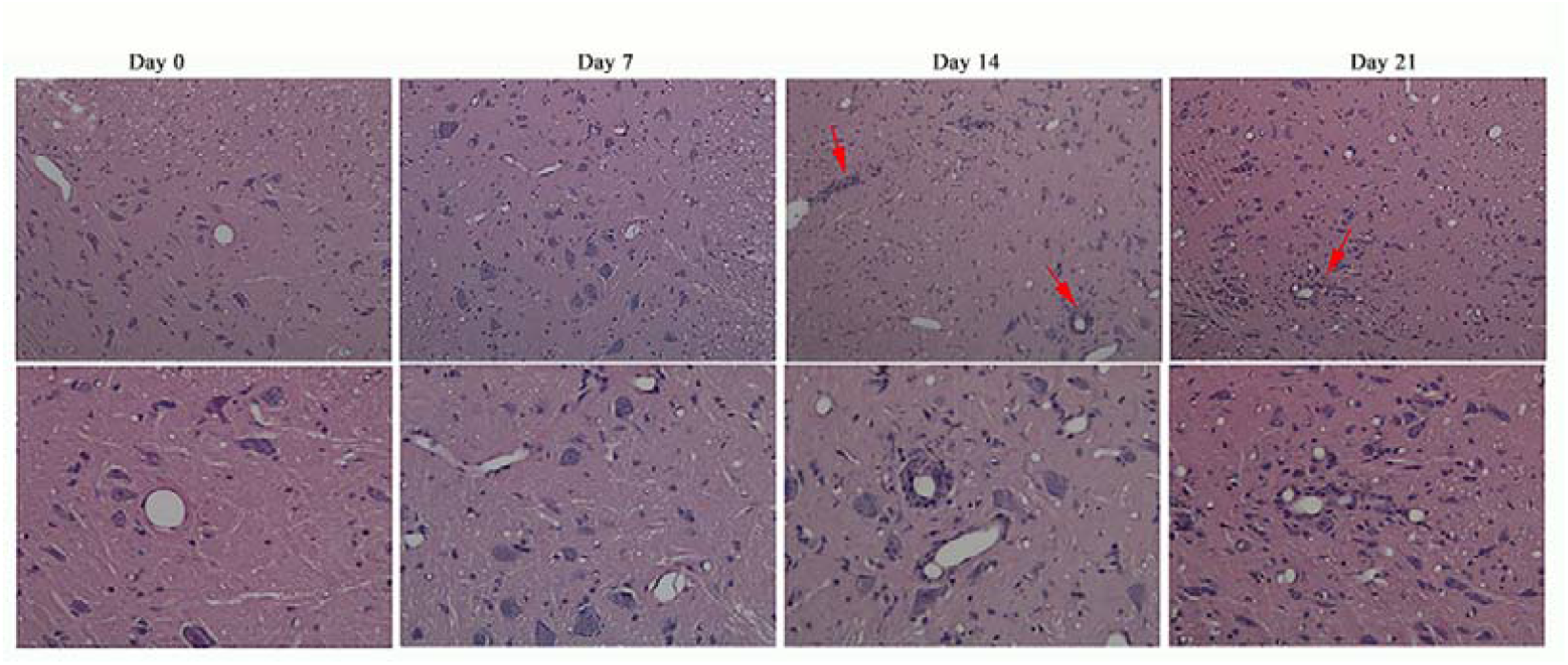
Histological characterization of the spinal cord in rats with EAE. A) Histological changes in the four stages in rats with EAE (H&E staining, 10×). B) Histological changes in the four stages in rats with EAE (H&E staining, 100×).

### Early intervention in EAE models by methylprednisolone

To explore the effect of early intervention, we established another EAE rat models injected with methylprednisolone at different time points to observe the incidence of the rats. For the early intervention group, all the EAE rats were received intraperitoneal injection of methylprednisolone on the 9th day after modeling for 5 consecutive days. Only one rat had onset of illness on day 11, the rest of the rats showed no obvious signs of nerve damage throughout the trial. For the late intervention group, the rat received intraperitoneal injection of methylprednisolone for 5 consecutive days after symptom onset. On day 10, two rats developed symptoms of nerve damage. All rats showed varying degrees of symptoms until day 12. For the EAE control group which injected with normal saline after onset, three rats had onset of illness on day 10. Other rats showed varying degrees of symptoms on day 11 and day 12. One rat in the late treatment group and two in the control group died during the experiment due to severe disease.

The changes in body weight and neural scores of the three groups are shown in Figure 4. From the 8th day, the body weight of the three groups decreased to varying degrees. After the nerve damage symptoms recovered on the 18th day, the body weight of all rats continued to increase. The experimental results showed that when EAE rats had onset of illness, whether or not methylprednisolone was used for treatment, the neurological damage symptoms of the rats gradually improved after the peak on day 15, which may be related to the spontaneous recovery of the model [16]. Compared the incidence of the three groups, the incidence in the early intervention group was only 1/13. The results indicate that early intervention can significantly reduce morbidity.

**Figure 4.**
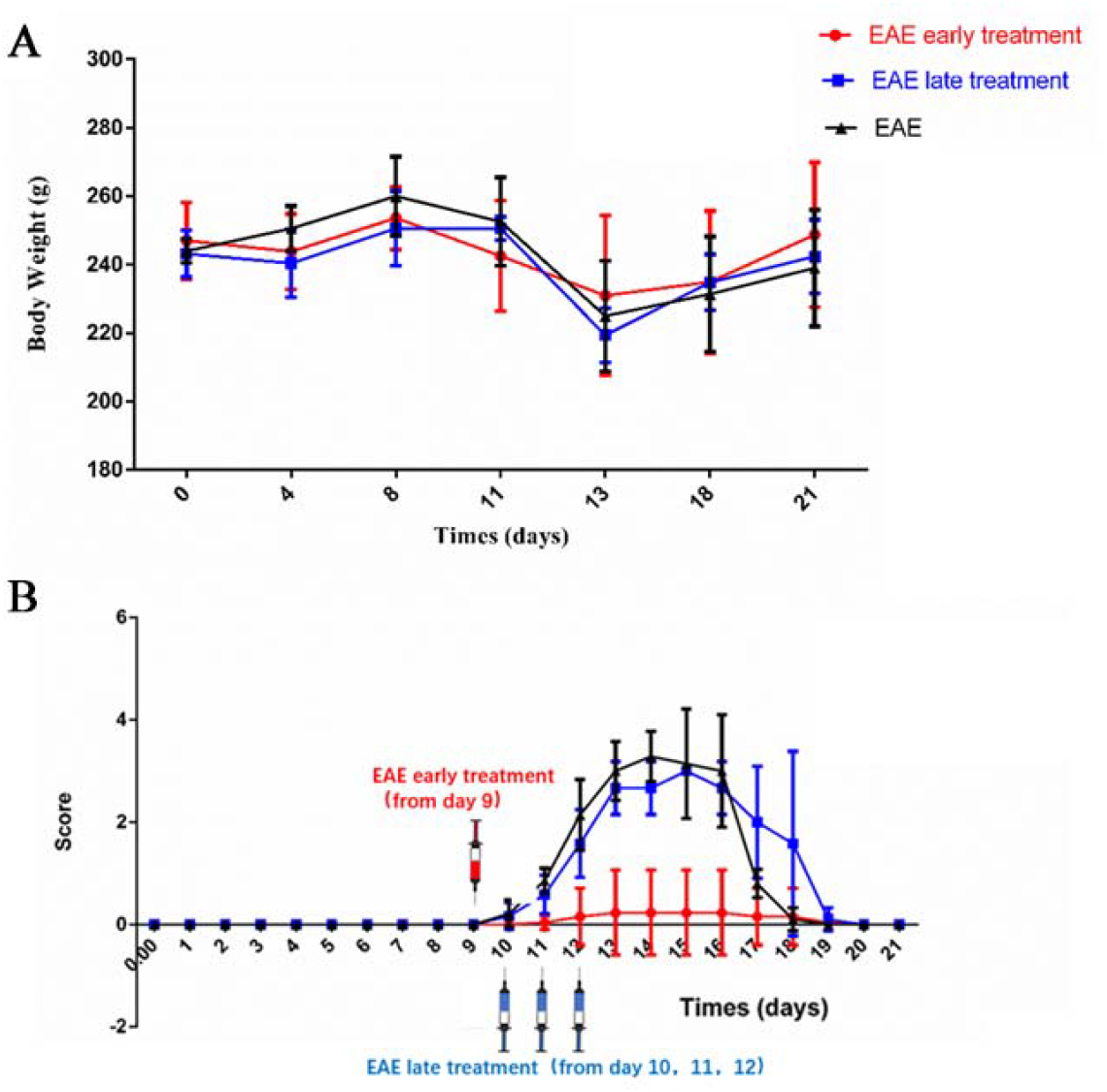
Changes in body weights and neurological damage score during the intervention experiment. A) Change in body weight of rats after modeling. B) The neurological injury score of rats after modeling.

### Early changes in urinary proteins in EAE models

As indicated above, on day 7, the clinical scores in the EAE group were “0” and were like the scores in the control group; no obvious histological changes were observed. Therefore, urine samples collected on day 7 after immunization were used for early biomarker detection. All identified proteins were quantitated by Scaffold Q+ software. The quantitative data are listed in Additional File 1. The altered proteins were defined based on the following parameters: p value < 0.05 (t-test) and fold change ratio > 1.5. Statistical analyses indicated that 25 proteins were significantly affected by MBP immunization (14 up- and 11 downregulated proteins). The proteins are listed in Table 1.

**Table 1.** The early dysregulated urinary proteins in EAE models.

| ID | Protein name | MW | Human orthologs | Fold Change E/N | p value | Type(s) | Reported in multiple sclerosis patients or animal model before |
| --- | --- | --- | --- | --- | --- | --- | --- |
| P01015 | angiotensinogen | 52 kDa | Yes | 2.16 | 2.66E-03 | growth factor | Serum/CSF[17] |
| P10760 | adenosylhomocysteinase | 48 kDa | Yes | 0.60 | 1.85E-02 | enzyme |  |
| Q8VI04 | asparaginase like 1 | 34 kDa | Yes | 1.78 | 4.00E-02 | enzyme |  |
| P47853 | biglycan | 42 kDa | Yes | 0.66 | 2.61E-02 | other |  |
| P08649 | complement C4B | 192 kDa | Yes | 1.66 | 1.08E-02 | peptidase | CSF[18] |
| Q62930 | complement C9 | 62 kDa | Yes | 1.73 | 2.27E-03 | other | CSF[18] |
| P10959 | carboxylesterase 1C | 60 kDa | No | 0.66 | 1.77E-02 | enzyme |  |
| P13635 | ceruloplasmin | 121 kDa | Yes | 1.52 | 7.50E-03 | enzyme | CSF[18] |
| Q9EQV9 | carboxypeptidase B2 | 49 kDa | Yes | 1.51 | 1.58E-03 | peptidase | CSF[19] |
| P12020 | cysteine-rich secretory protein 1 | 28 kDa | No | 3.18 | 1.28E-06 | other |  |
| P55054 | fatty acid binding protein 9 | 15 kDa | Yes | 6.17 | 3.32E-02 | transporter | Serum[20] |
| P20059 | hemopexin | 51 kDa | Yes | 1.82 | 2.29E-03 | transporter | CSF[18] |
| P01048 | T-kininogen 1 | 48 kDa | Yes | 3.18 | 3.87E-03 | other | CSF[18] |
| Q9QZ76 | myoglobin | 17 kDa | Yes | 0.56 | 1.76E-02 | transporter |  |
| O88989 | malate dehydrogenase 1 | 36 kDa | Yes | 0.59 | 7.12E-03 | enzyme |  |
| P70490 | milk fat globule-EGF factor 8 protein | 47 kDa | Yes | 1.50 | 1.79E-02 | other |  |
| O88766 | matrix metalloproteinase 8 | 53 kDa | Yes | 2.30 | 6.20E-03 | peptidase | Tissue/plasma[21] |
| P02631 | oncomodulin | 12 kDa | Yes | 0.61 | 2.64E-03 | other | Tissue[22] |
| P02764 | orosomucoid 1 | 24 kDa | Yes | 2.32 | 4.77E-02 | other | CSF[18] |
| P10111 | peptidylprolyl isomerase A | 18 kDa | Yes | 0.56 | 1.77E-02 | enzyme |  |
| Q62714 | defensin RatNP-3 precursor | 10 kDa | Yes | 2.73 | 1.50E-02 | other |  |
| P50116 | S100 calcium binding protein A9 | 13 kDa | Yes | 0.37 | 1.60E-02 | other | Plasma[23] |
| P07092 | serpin family E member 2 | 44 kDa | Yes | 0.51 | 8.29E-05 | other | CSF[24] |
| P09656 | serine protease inhibitor Kazal-type 1-like | 9 kDa | Yes | 0.65 | 3.35E-02 | other |  |
| P11232 | thioredoxin | 12 kDa | Yes | 0.60 | 2.08E-03 | enzyme | Serum[25] |

### Canonical pathway and proteomic interactomes of the changed proteins

The canonical pathway that the changed proteins involved were generated by IPA software. The most significant pathways are acute phase response signaling, LXR/RXR activation, FXR/RXR activation, complement system and asparagine degradation (Figure 5A). The other pathways are mostly related to the acute response except for asparagine pathway. Though its role in multiple sclerosis is unclear, the asparagine may have roles in proper immune functioning and asparagine biosynthesis may be one of the main canonical pathways involved in multiple sclerosis [26]. The inhibition of asparagine endopeptidase greatly enhanced presentation of the myelin basic protein [27], which is a known key point of multiple sclerosis. As a result, the asparagine degradation pathway may play important roles in the disease onset and progression.

**Figure 5.**
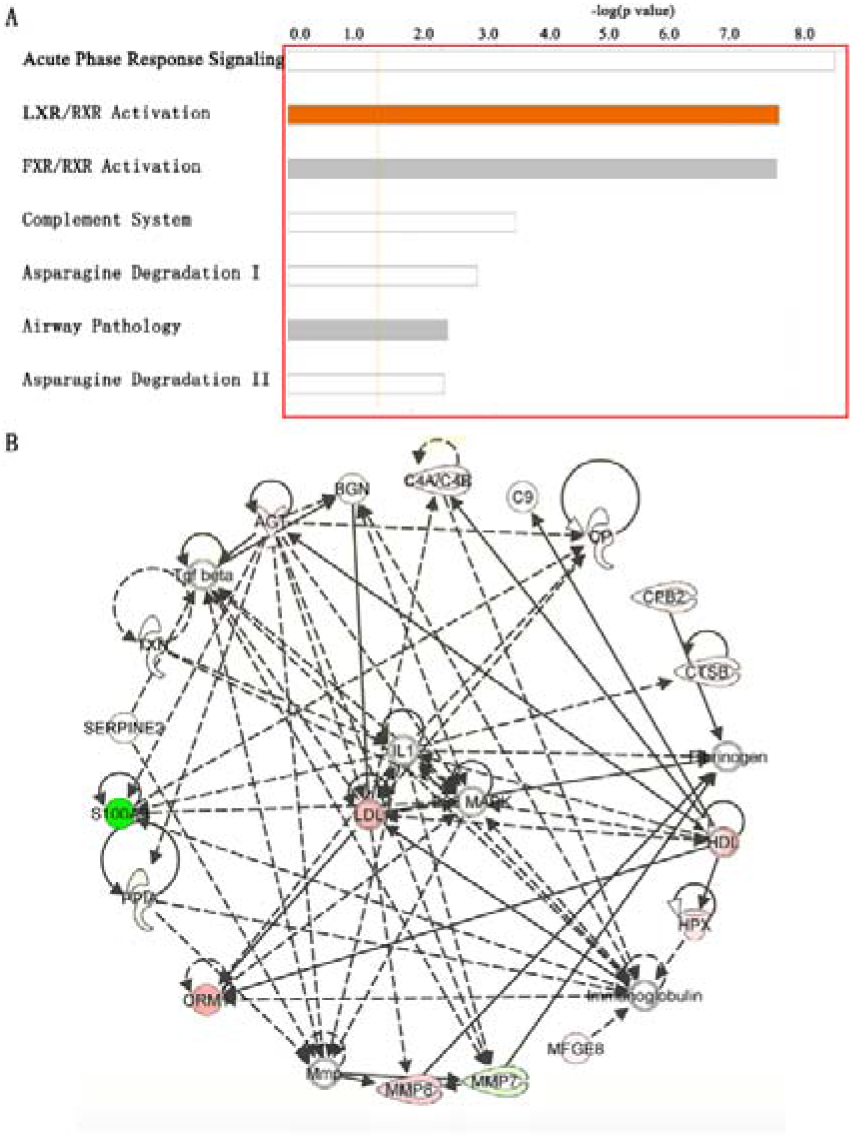
Functional analysis of changed proteins by IPA software. A) The top canonical pathways generated from the software. B) Network related to neurological functions annotated by IPA.

To investigate the analytical underpinning of this urinary proteomics study, the proteomic interactome of altered proteins after MBP immunization was determined. The IPA software highlights interactions among specific molecules and demonstrates how they might work together at the molecular level. Thus, these networks might represent an efficient and unified interactions among the deregulated proteins and highlight potential causal molecules that may be used as targets for preventing multiple sclerosis progression. As shown in Figure 5B, we uploaded 25 deregulated proteins into IPA and determined their interactions with network proteins associated with “neurological diseases”. A total of 13 deregulated molecules were identified in this functional interaction. Specific peptidases (complement C4, carboxypeptidase B and matrix metallopeptidase 8), enzymes (ceruloplasmin, peptidylprolyl isomerase A and thioredoxin), transporter (hemopexin), growth factor (angiotensinogen) and others (biglycan, complement C9, orosomucoid 1, S100 calcium binding protein A9 and serpin family E member 2) were identified as being altered in MBP-induced EAE. Within these networks, highly interconnected hub molecules are more likely to have important biological functions. The hub proteins identified in the neurological disease networks, namely, IL-1, LDL and P38 MAPK, are in the extracellular space, plasma membrane and cytoplasm, respectively. The upregulation of P38 MAPK is closely related to 4-1BB signaling in T cells and is consistent with the induction of EAE, because EAE is initiated by immunization with autoantigens presented to MHC class II-restricted CD4+ T helper cells [9]. Additionally, the inhibition of active mouse p38 MAPK in CD4+ T cells was shown to decrease the severity of EAE in mice [28]. IL-1 is one of the commonly used inflammatory factors. Although it is rarely detected in the normal brain, IL-1 is significantly upregulated and plays a central role in neuroinflammation, especially under neurodegenerative conditions [29]. Therefore, the proteins dysregulated in response to MBP immunization include interactors that preferentially function in CD4+ T cells, neuroinflammation and lipid metabolism and hence may affect neurological functions.

### Human homologs of the altered urinary proteins

Altered proteins that are homologous to human proteins are potentially useful in clinical practice and may be candidate biomarkers of multiple sclerosis. When we imported the 25 altered proteins into the InParanoid database [30] to search for human homologs, 23 of the 25 proteins were found to have human counterparts (Table 1). Because these 23 proteins may be useful for clinical practice, we will discuss these proteins that have human homologs in the following part of the manuscript.

Among the twenty-three urinary proteins that were affected and had a relatively significant fold change compared with the control group, fourteen were reported to be differently expressed in the serum/ cerebrospinal fluid (CSF)/brain tissues of multiple sclerosis (Table 1). Among them, thioredoxin was thought to be dysregulated in multiple sclerosis patients exposed and non-exposed to treatment [25] and serpin family had a significant and reproducible correlation with multiple sclerosis severity [24]. These findings may indicate the accuracy and validity of the MS analysis and demonstrate urine as a good source of biomarkers for multiple sclerosis.

The dysregulated proteins in urine are consistent with the previous studies in CSF and plasma. For example, both kininogen (a precursor for kinin) and complement component 9 are mediators of inflammation and play important roles in response to inflammatory injury. Elevated levels of kininogen and complement 9 in the CSF have been reported in rats that have EAE [18]. Additionally, expression of the kinin B1 receptor mRNA on peripheral blood mononuclear cells can serve as an index of disease activity in multiple sclerosis [31]. Therefore, in EAE or multiple sclerosis, the expression of kininogen is upregulated in the plasma, CSF, and urine. Protease family members, including metalloproteases, serine proteases, and cysteine proteases, can be markers of disease activity in multiple sclerosis [21]. Serum fatty acid binding protein (FABP) is thought to distinguish subtypes of multiple sclerosis, because it is expressed at the highest level in secondary progressive multiple sclerosis and increased during early stages of pediatric-onset multiple sclerosis [20]. In the current study, urinary FABP 9 level was also increased in the early stages of disease. Oncomodulin, a factor produced by macrophages, promotes axon growth in neurons and is an indicator of central nervous system injury [22], including multiple sclerosis. In the early stages of EAE, the levels of oncomodulin are reduced, which may partly be due to axonal injury. The neutrophil collagenase (MMP-8), a metalloprotease, has been shown to increase in the central nervous system in response to EAE and is correlated with symptom severity [21]. Compared with MMP-8 in the central nervous system, MMP-8 in urine is upregulated by as much as 1.5-fold during the early stages of EAE and can further increase with disease progression. Angiotensinogen is involved in maintaining blood pressure and in the pathogenesis of essential hypertension and preeclampsia. Interestingly, the upregulation of serum angiotensin-converting enzymes is related to disease activity in longitudinal analysis [32], while reduced levels of intrathecal angiotensin II in the CSF are indicators of neural damage and repair processes in multiple sclerosis [33]. Consistent with plasma angiotensinogen, urinary angiotensinogen was also increased after immunization in rats that have EAE.

As most of the significantly altered proteins in this study are related to multiple sclerosis; this finding may indicate the accuracy of MS analysis. This study may further illustrate the clinical value of urinary proteome in studying the disease in central nervous system. These differential proteins that have not been confirmed in the literature still need to be further evaluated for their diagnostic value. And more clinical samples are required to assess the significance of differential proteins in this experiment.

## Conclusion

In this study, early candidate urinary biomarkers of multiple sclerosis were identified in a rat model of EAE before histological changes and clinical symptom onset. More than half of the differentially regulated proteins were identified as participants in neurological functions and reported to be related to the multiple sclerosis. Additionally, intervention at the early time points when urinary protein has changed can effectively reduce the morbidity of EAE models. In conclusion, this study showed that urine may be a good source of multiple sclerosis early biomarkers. These findings may provide important information for early diagnosis and intervention of multiple sclerosis in the future.

